# Real-Time, Direct Classification of Nanopore Signals with SquiggleNet

**DOI:** 10.1101/2021.01.15.426907

**Authors:** Yuwei Bao, Jack Wadden, John R. Erb-Downward, Piyush Ranjan, Weichen Zhou, Torrin L. McDonald, Ryan E. Mills, Alan P. Boyle, Robert P. Dickson, David Blaauw, Joshua D. Welch

**Affiliations:** Department of Computer Science and Engineering, University of Michigan, 48109 Ann Arbor, Michigan, USA; Department of Computational Medicine and Bioinformatics, University of Michigan, 48109 Ann Arbor, Michigan, USA; Department of Electrical and Computer Engineering, University of Michigan, 48109 Ann Arbor, Michigan, USA; Division of Pulmonary and Critical Care Medicine, Department of Internal Medicine, University of Michigan Medical School, 48109 Ann Arbor, Michigan, USA; Department of Human Genetics, University of Michigan Medical, 48109 Ann Arbor, Michigan, USA; Department of Microbiology and Immunology, University of Michigan Medical School, 48109 Ann Arbor, Michigan, USA; Michigan Center for Integrative Research in Critical Care, 48109 Ann Arbor, Michigan, USA

**Keywords:** Deep Learning, Read-Until, Oxford Nanopore, Raw signal, Real-time

## Abstract

Oxford Nanopore sequencers provide results in real time as DNA passes through a nanopore and can eject a molecule after it has been partly sequenced. However, the computational challenge of deciding whether to keep or reject a molecule in real time has limited the application of this capability. We present SquiggleNet, the first deep learning model that can classify nanopore reads directly from their electrical signals. SquiggleNet operates faster than the DNA passes through the pore, allowing real-time classification and read ejection. When given the amount of sequencing data generated in one second, the classifier achieves significantly higher accuracy than base calling followed by sequence alignment. Our approach is also faster and requires an order of magnitude less memory than approaches based on alignment. SquiggleNet distinguished human from bacterial DNA with over 90% accuracy, generalized to unseen species, identified bacterial species in a human respiratory meta genome sample, and accurately classified sequences containing human long interspersed repeat elements.

## Introduction

Oxford Nanopore sequencers, such as MinION or PromethION, determine the nucleotide sequence of a DNA or RNA molecule by measuring changes in electrical current (called “squiggles”) as the molecule translocates through a protein nanopore. This approach is fundamentally different from the widely-used Illumina platform and provides several benefits: the MinION is small, fast, and portable, making it ideal for rapid diagnostics and field work. Because it does not rely upon synchronized nucleotide addition (the heart of the Illumina sequencing-by-synthesis technology), MinION also produces much longer reads. To our knowledge, the longest published MinION read is around 2Mbp [1], though even longer reads have been reported anecdotally. The changes in electrical current induced by a DNA or RNA molecule depend on the specific chemical properties of the nucleotides, including secondary structure interactions and epigenomic modifications such as methylation. Additionally, the nanopore sequencer can stream the squiggle data to a computer in real time.

The nanopore sequencer can also eject a partially sequenced molecule, a capability referred to as “Read-Until”. In principle, this enables targeted sequencing without the need for biochemical enrichment. The Read-Until capability allows selective sequencing of molecules by reversing the voltage across individually selected nanopores, ejecting the unwanted molecules. The unoccupied nanopores can then sequence different molecules of interest.

Such computational enrichment of target sequences holds great promise for clinical diagnostics and field research, but realizing this potential requires fast and accurate approaches for identifying molecules of interest. For example, identifying pathogenic DNA in a patient lung fluid sample requires bypassing human DNA – which often represents *>* 99% of the sequences – to find the pathogen sequences. Biochemical methods for target sequence enrichment, such as PCR [2–5], hybrid capture [6], or CRISPR/Cas9 enrichment [7, 8] require much more time, expertise, and equipment. In contrast, a computational approach to enriching target sequences provides clear savings of time, labor, and cost.

Previous computational approaches for this problem include: a) perform standard base calling followed by sequence alignment as in [9], and b) perform rough base calling to identify and align *k*-mers [10]. The first approach requires significant computing resources – such as a graphics processing unit (GPU) and a large genome index database for the sequence aligner. The second approach also requires a large genome index and multiple CPU cores and can map only non-repetitive references smaller than ∼100Mbp. Both approaches are based on sequence alignment and thus are limited by sequencing errors, their reliance on genome indexes, and their inability to capture non-sequence information such as DNA methylation.

To address these limitations, we developed SquiggleNet, the first deep-learning based approach for classifying DNA sequences directly from electrical signals. SquiggleNet is fast, accurate, memory-efficient, and robust to unknown species. It requires only 3000 signals – less than the amount of data generated in one second of sequencing – to classify the species of a DNA molecule with over 90% accuracy, significantly higher than the best alignment-based methods. The model requires only 304 KB of RAM and no external reference database. SquiggleNet is faster than or on par with the competitors and can run in real time on a single core of a standard laptop. When tested on a human respiratory metagenome sample with a majority of unseen species, our approach achieves > 90% overall accuracy.

## Results

### SquiggleNet: A Convolutional Neural Network for Classifying Nanopore Signals

SquiggleNet is a deep neural network that classifies molecules of interest based on statistical patterns in nanopore conductivity, which are often hard for humans to identify by eye, automatically extracted from the input data. The overall workflow for using SquiggleNet to enrich sequences of interest is shown in Figure 1a. The network is first trained to recognize certain classes of sequences, such as human vs. bacterial DNA, using labeled examples. Then, as the nanopore sequencer generates raw electrical signals from a new and unseen sample, SquiggleNet rapidly classifies each molecule to determine whether it is a sequence of interest. Molecules not of interest are ejected from the nanopore, freeing the pore to sequence a different molecule. In contrast, targeted molecules are sequenced to full length and used for downstream analysis.

**Figure 1:**
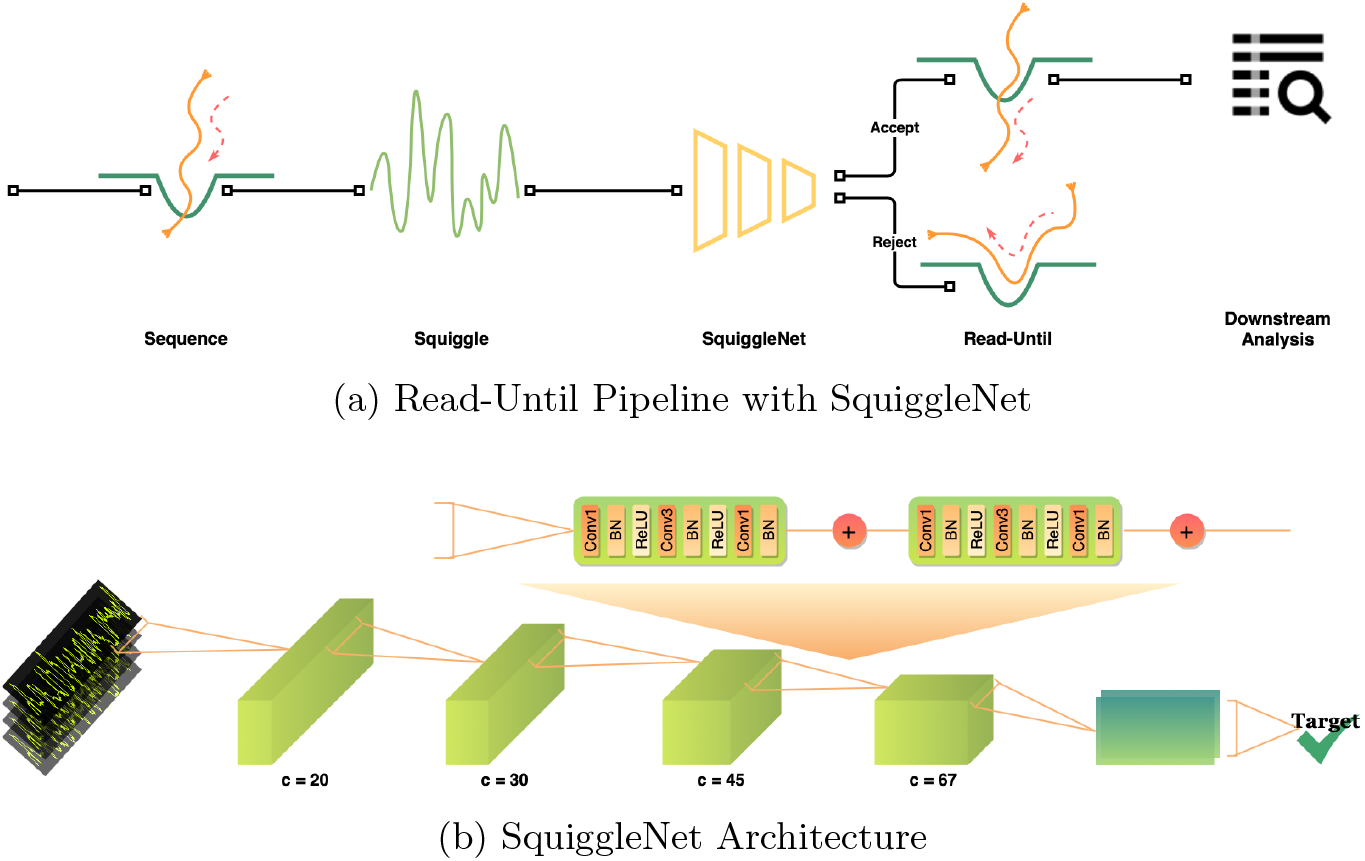
Read-Until Pipeline Overview: **(a)** A DNA molecule translocates through a nanopore, generating electric signals (squiggles). SquiggleNet rapidly classifies the molecule to determine whether it is a sequence of interest. If the molecule is accepted by the classifier, it is sequenced to full length. Otherwise, the molecule is ejected from the pore, freeing the pore to sequence another molecule. **(b)** SquiggleNet employs 1D ResNet bottleneck blocks with increasing numbers of filters. We perform average pooling after the last convolutional block and use a final fully connected layer.

SquiggleNet (Figure 1b) employs a convolutional architecture, using residual blocks modified from ResNet [11] to perform one-dimensional (time-domain) convolution over squiggles. The architecture consists of four blocks with increasing numbers of channels; each block includes two 1-D ResNet Bottleneck units. Mean pooling followed by a fully-connected layer with softmax activation allows SquiggleNet to classify sequences based on the convolutional filters in the last ResNet block. The final output is a conditional probability on the sequence labels, which is then used to make the final class prediction. We experimented with several other approaches, including a recurrent neural network (RNN) with long short-term memory (LSTM) blocks; gated recurrent units (GRUs); other types of convolutional blocks; a combination of RNN and convolution; different convolutional window sizes; and differing model hyperparameters. However, we found that approaches based on convolution outperformed models using LSTM blocks, suggesting that local features are sufficient for this problem, and long-range time-dependent relationships do not add much information. Convolutional architectures without LSTM blocks are also faster to train. Our final architecture gave the best classification accuracy of any approach we tried and could not be made significantly smaller without sacrificing performance. Additional details about the model architecture and hyperparameter choices can be found in the Method section and Supplement.

### SquiggleNet Accurately Classifies Species Directly from Squiggles

To test the performance of SquiggleNet, we generated four experimental datasets containing a mixture of human and bacterial DNA. The first dataset, HeLa&Zymo, contains 8 bacterial species from the Zymo mixture [12] and HeLa cells. The species labels were obtained through Minimap2 [13] alignment. The other three datasets (Human&Zymo b12, Human&Zymo b34, and Human&Zymo b56) contain a mixture of human GM12878 DNA and DNA from Zymo High Molecular Weight mixture with 7 bacterial species [14]. To avoid systematic error from the alignment algorithms, we obtained reliable ground-truth species labels for these three datasets by attaching a nucleotide sequence barcode to each DNA molecule indicating whether the molecule is from human or bacteria. Note that our direct biochemical labeling strategy allows us to independently assess the accuracy of species determination from base calling followed by read alignment; this is important for our application, since we expect that SquiggleNet may be able to outperform purely sequence-based approaches by leveraging other information from the electrical signals. Further dataset details can be found in the Method section and Supplement.

We trained SquiggleNet using more than two million reads from the first dataset (HeLa&Zymo), which contains equal proportions of HeLa and bacterial sequences. We used 3000 signals from each read, the equivalent of about 300 nucleotides. We discarded the first 1500 signals of each read (an overestimation of the adapter length), to remove potential pore noise and adapter sequences, which could confound training. Thus SquiggleNet requires a total of 4500 signals, which is equivalent to about 1 second of sequencing time. (The exact time and number of nucleotides depends on the translocation speed, which varies per pore and molecule over the course of the sequencing experiment.) However, using this exact amount of signal is not crucial; we verified that using fewer signals did not significantly change the results (see below). Our best-performing model was trained on the HeLa&Zymo dataset, which contains the largest number of sequenced reads. This dataset also lacks species-specific barcodes, and we were careful to remove the sequencing adapters and species barcodes before extracting the 3000 signals used for classification (Methods). Thus, there is no way that the classifier could “cheat” by using the barcodes to classify the species. When we instead trained the model on the Human&Zymo datasets and tested on HeLa&Zymo, the model accuracy was nearly identical but slightly lower, possibly due to the smaller number of training samples (see Supplement and Fig. 9).

Overall, the model classifies each molecule as bacterial or human with over 90% accuracy across different test datasets using only 3000 signals per read (see Figure 2). The classifier generalizes well to different lab preparations, flow cells and proportions of species. For the first three datasets (HeLa&Zymo, Human&Zymo b12, Human&Zymo b34), the true positive rates (TPR, also Recall) and the true negative rates (TNR) are all above or around 90%. The precision and AUROC scores are all about 90% as well. Even for samples with significantly more human than bacterial DNA (Human&Zymo b56 and Respiratory Metagenome), the accuracy and recall both remain high.

**Figure 2:**
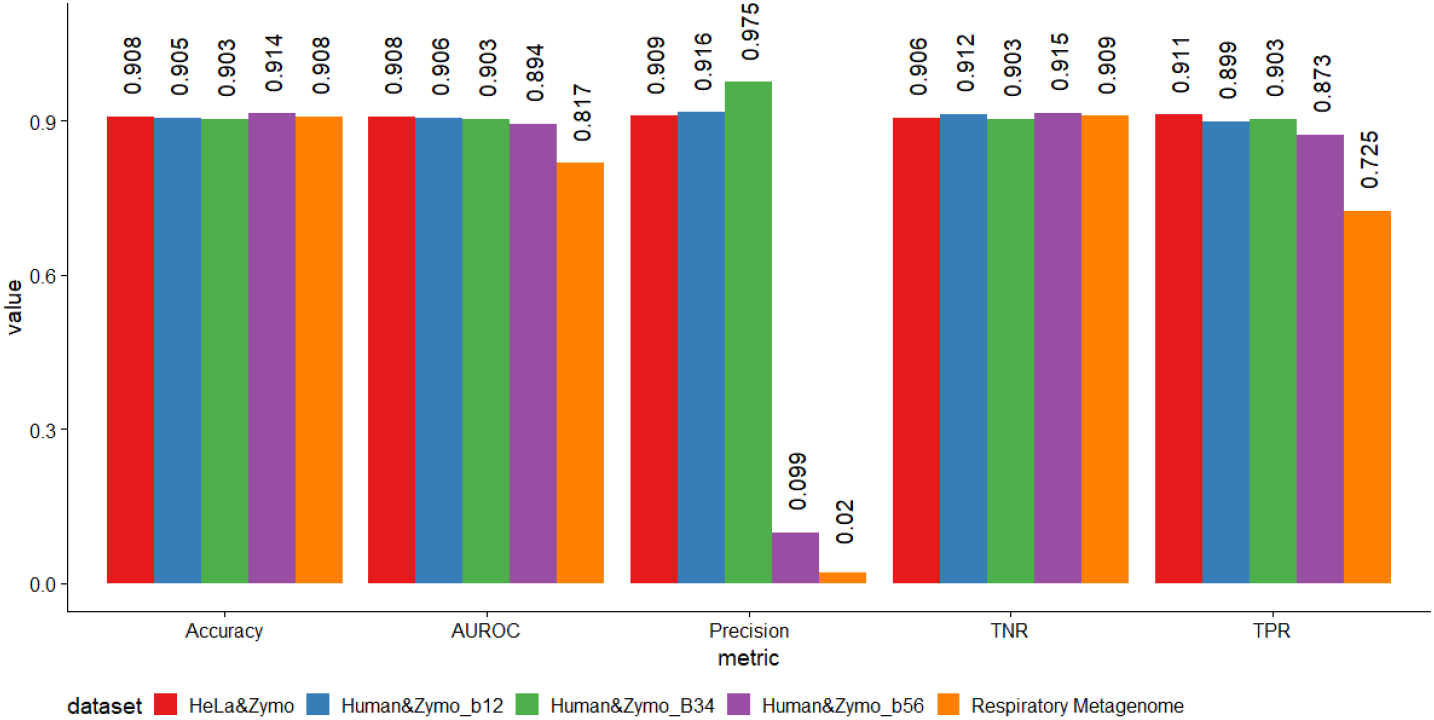
Overall Performance across five test Datasets: Accuracy, True Positive Rate(TPR, RECALL), True Negative Rate(TNR), Precision, and the AUROC score of the model trained on the HeLa&Zymo training set, and tested on five test sets with bacterial sequences as the target.

We used the method of integrated gradients (IG) [15] to investigate the features influencing SquiggleNet’s classification decisions. The IG method computes the amount of gradient change for each corresponding input, and by doing so, offers interpretation on which part of the input contributes the most to the model’s decision. Inspecting these IG results (Fig. 10) shows that SquiggleNet predictions are most strongly influenced by positions where the signal changes direction, changes by a large amount, and/or changes from one nucleotide to another. This suggests that SquiggleNet has learned filters related to the nucleotide composition of the signal and uses the results to make classification decisions. Further details are contained in the Supplement.

To demonstrate that the results are robust to the amount of signal removed from the beginning of the read at test time, we also tested our pre-trained model on the Human&Zymo b34 dataset with only the first 1000 signals per read removed. We chose the number 1000 because this is a closer estimation of the adapter length [16], and at test time, we would like to make the decision as soon as possible to enable real-time read selection. When testing the model on sequences with the first 1000 signals removed, the results were nearly identical to those obtained from conservatively removing 1500 signals: 89.35% accuracy, 90% true positive rate, and 86.9% true negative rate. Thus it appears that, as long as the initial pore noise, adaptors, and barcodes are removed from the training sequences, the model is able to make an accurate and fast decision at test time. This robustness also allows flexibility if, for example, different sequencing datasets use sequencing adapters of different lengths.

Remarkably, we find that SquiggleNet achieves significantly higher accuracy from 3000 signals than base calling followed by sequence alignment using the same amount of signal. This result gives crucial context for interpreting the accuracy of our model and suggests that the convolutional filters may detect some non-sequence features that help with species classification, such as chemical modification of nucleotides by methylation. Indeed, we found that the bacterial and human DNA sequences in our dataset show significant methylation differences, with significantly more methylated cytosines in human sequences and significantly more methylated adenines in bacterial sequences (see Supplement and Table 4 for details).

For the Human&Zymob 56 dataset, the target to non-target sequence ratio is 1 to 99. The overall accuracy, TNR, and AUROC score are around 90%. The TPR (Recall) is closely following, above 87%. The precision, however, is about 1*/*10 of the other cases. This is due to the extremely low concentration of the target sequences (Zymo bacterial species), and the precision calculation is diluted by the overwhelming number of false positive reads. Nevertheless, this is acceptable, since we want to preserve as many targeted reads as possible (high recall) due to the low target read concentration. All true positives and false positives will be sequenced to full length, and thus can be further processed in the downstream analysis. Considering that the Human&Zymo b56 dataset has 99× more non-targeted reads than targeted, whereas only ∼10× more reads were falsely identified as positive compared to the HeLa&Zymo dataset, this model demonstrated strong ability to filter out non-targeted reads, and has high potential to improve throughput (see below). Overall, the model that was trained on only the HeLa&Zymo dataset yields high performance across different testing datasets, highlighting the robustness of the model.

Interestingly, SquiggleNet performance varies systematically across bacterial taxa. The network classifies human vs. bacterial DNA with 90% accuracy, but some bacterial species are easier to distinguish from human sequences than others. The eight bacterial species in the Zymo mixture are related according to the taxonomy tree shown in Figure 3. The top three species–*Pseudomonas aeruginosa* (Pse), *Salmonella enterica* (Sal), and *Escherichia coli* (Esc)–are gram-negative bacteria and are most easy to identify, while the bottom five species are gram-positive bacteria and are harder to distinguish from human DNA. It is not clear what specific features of the gram-negative bacteria make them easier to identify, but this behavior may be related to species differences in GC-content or the amount of methylation.

**Figure 3:**
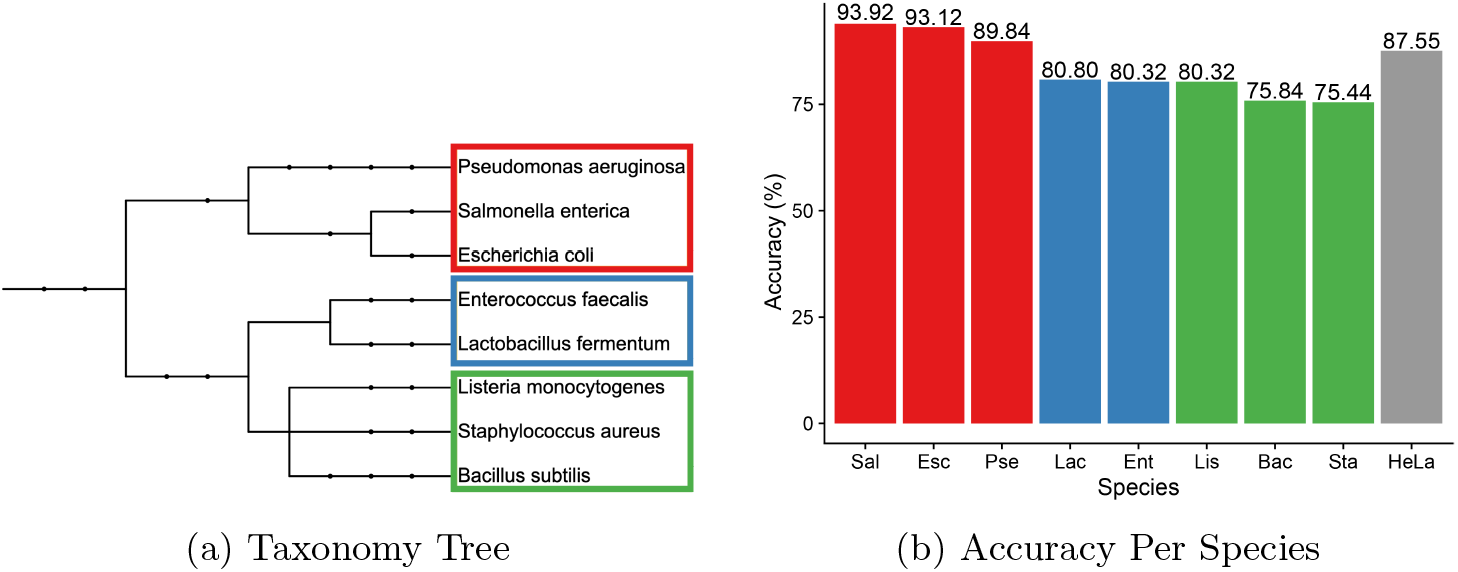
Taxonomy Tree and Accuracy Per Species: Taxonomy tree for the eight species in our dataset grouped in color and their corresponding accuracy breakdown per species. The accuracy for distinguishing bacterial sequences from human was highest for the red branch, intermediate for the blue group, and lowest for the green group.

### SquiggleNet Identifies Species Not Seen During Training

In real-world applications, samples may contain species whose genomes are not in the training samples. We thus investigated whether the model can identify unseen species. To do this, we performed a leave-one-out analysis, removing each of the bacterial species separately during training, then putting it back during testing to challenge SquiggleNet’s generalization ability. For the held-out species comparisons, we used 400k and 20k reads from the HeLa&Zymo dataset for training and testing, respectively.

During each training run, we removed one of the eight Zymo bacterial species from the training dataset. We then compared the test accuracy from the classifier trained on seven bacterial species plus human with the performance of the same model on two different testing sets containing the eighth held-out species. The dataset we call Test-Uniform/HeLa includes all eight species (including the one held out during training), evenly distributed, and balanced to contain equal numbers of HeLa and bacterial molecules. The dataset we refer to as Test-One/HeLa includes only the single held-out species and HeLa, in equal proportions.

The unknown species identification results can be found in Figure 4. The red bars are the test accuracy results without held-out species. The left-most column is the performance of a training run with all 8 bacterial species as a reference for cross-testing run performance comparison.

**Figure 4:**
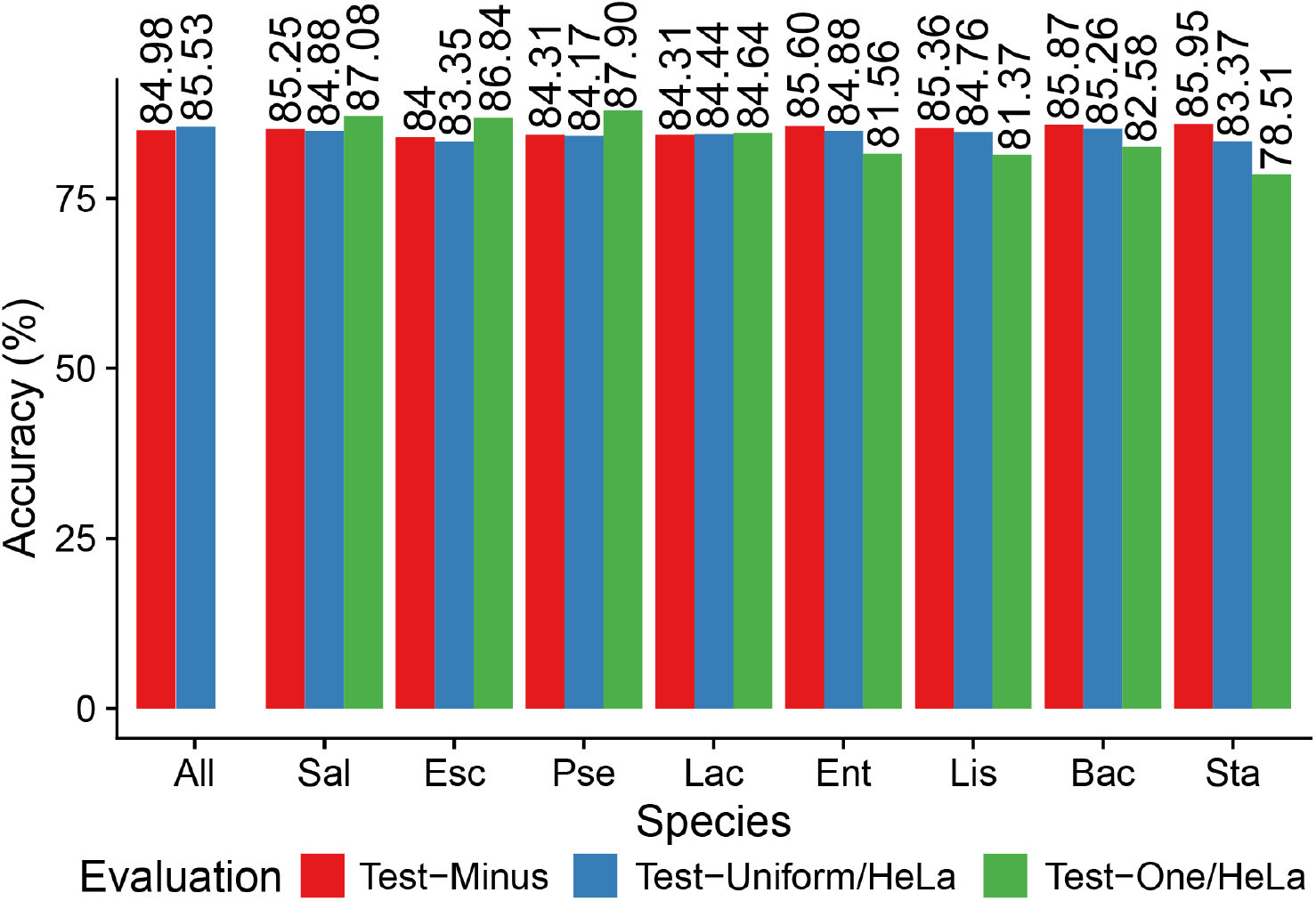
Performance of SquiggleNet on Unseen Species: Each column (except “All”) is a model trained on a Zymo/HeLa 1:1 mix without the held-out species. For each species, the red bar shows the test accuracy on all species minus the held-out species; this number provides a baseline against which to compare performance on the held-out species. Blue bars show the accuracy of each trained model on Test-Uniform/HeLa, a test set with all eight Zymo bacterial species included and HeLa in a 1:1 ratio. Green bars show the accuracy of each model on Test-One/HeLa, a test set with only the single unseen species and HeLa in a 1:1 ratio.

Across different runs, the test accuracies, not including held-out species, are around 84%-86%. For each Test-Uniform/HeLa experiment, accuracy of classifying the held-out species was ∼83%-85%, only about 1% lower compared to when the species was seen during training. This shows that the model was able to accurately identify sequences from bacterial species that were not seen during training. For the Test-One/HeLa experiment, the test performance is more influenced by the taxonomic position of the held-out species. Since the testing datasets only include human DNA and the one species that was held out, we expected performance to drop even more than the previous Test-Uniform/HeLa experiment. However, the test accuracies of the first three gram-negative bacterial species, *Pseudomonas aeruginosa* (Pse), *Salmonella enterica* (Sal), and *Escherichia coli* (Esc) actually increased by ∼1%-4% compared to their validation accuracies. The remaining four gram-positive species had a minor performance increase or drop within 4%. *Staphylococcus aureus* had the largest performance drop among all, but the accuracy was still above 75%. In summary, these two sets of experiments show that even when one species was not seen during training time, SquiggleNet was still able to identify it with high confidence.

### SquiggleNet Identifies Bacterial DNA in a Human Respiratory Metagenome Sample

To further test the generalizability and practicality of SquiggleNet, we tested the best performing model (trained on the Hela&Zymo dataset) on a dataset collected from several clinical human samples. We collected the data following the procedures in [17]. The ground truth labels were obtained using our previously published read alignment pipeline[18]. The dataset includes 324526 human reads and 341 bacteria and other (less than 0.6%), a human:bacterial ratio of 951:1. Some of the dominant bacteria groups include *Prevotella* (29%), *Neisseria* (20%), and *Rothia* (11%). However, less than 3% of the bacterial species overlap with the training dataset. The full taxonomic composition can be found in Figure 5 and [17].

**Figure 5:**
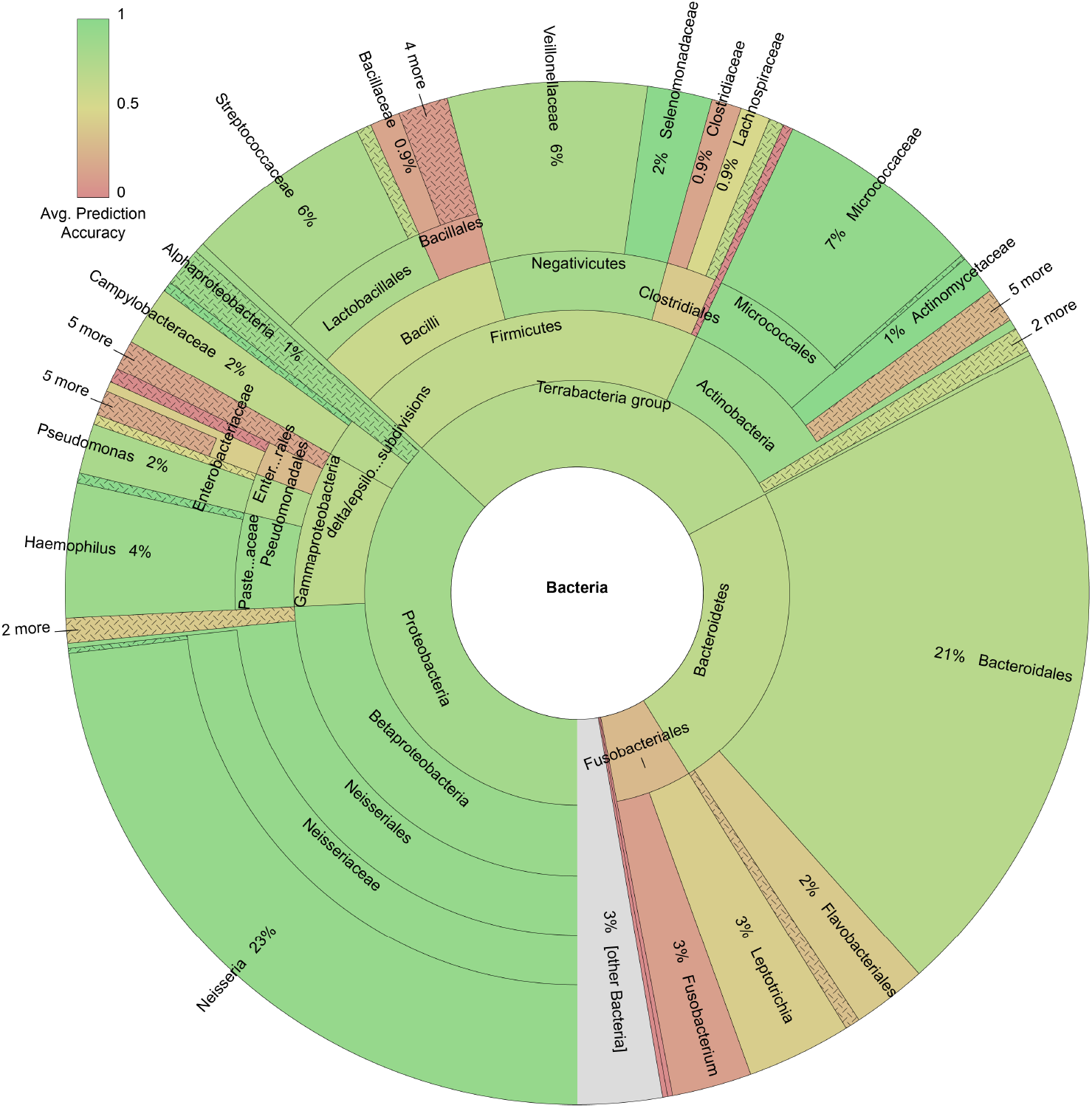
SquiggleNet Accuracy by Species in Human Respiratory Metagenome Sample[19]: Some of the dominant bacterial groups include *Neisseria* (23%), *Bacteriodales* (21%), and *Firmicutes* (20%). Less than 3% of the bacterial species overlap with the training dataset.

Even though the model was trained on dataset Hela&Zymo, it achieved 90.8% overall accuracy in the Respiratory Metagenome dataset, 72.5% true positive rate, and 90.9% true negative rate (Figure 2). The AUROC score is 0.817. The precision is about 1*/*5 of that in dataset Human&Zymo b56. As with the unbalanced Human&Zymo dataset, the precision is diluted by the extremely low concentration of bacteria, but the model still achieves high recall–which is critical to retrieve all the bacterial reads for downstream analysis.

The Zymo community of the dataset on which the model was trained has very little overlap (*<* 3%) with the bacterial species found in the Respiratory Metagenome dataset. The genome information in this testing dataset was mostly unseen and unknown for the trained model. However, it still achieved a True Positive Rate of 72.5%. This shows that SquiggleNet is able to extract common bacterial genome features and distinguish them from the human genome sequencing raw signals. The generalizability of SquiggleNet significantly increases the potential applications of our method. As shown in Figure 5, different species were classified with different accuracy. The model is therefore, expected to be even more accurate if it can be fine-tuned in a dataset with closer range of species.

### SquiggleNet Is More Accurate and Efficient than Previous Approaches

We next compared the performance and efficiency of SquiggleNet against the current state-of-the-art methods: Guppy+Minimap2 and UNCALLED. This experiment was conducted on dataset Human&Zymo b34 with 1:4 Human and Zymo mix. All the analysis was done on a single-usage Intel(R) Xeon(R) CPU E5-2697 v3 @ 2.60GHz machine with a single TITAN Xp GPU.

We benchmarked the running time required to classify 712,000 reads (178 fast5 files with 4000 sequences each and 3000 signals per read, adapters and barcodes removed). SquiggleNet took 806.74 seconds (13 minutes 27 seconds) to finish processing all on GPU (Figure 6). When tested on a 3.5 GHz Dual-Core Intel Core i7 Macbook Pro, SquiggleNet finished processing all the files in 2630.58 seconds (43 mins 51 seconds). With the Guppy+Minimap2 method [9], sequences were base called using Guppy [20] with 4 callers and 4 runners/GPU, and we used 32 threads for sequence alignment with Minimap2 [13]. It took 742.602 seconds (12min 23s) for Guppy to finish basecalling 3000 signals and another 25.673 seconds for Minimap2 to finish the alignment, about the same amount of time as SquiggleNet. The accuracy of Guppy+Minimap2, however, was 79%, more than 10% lower than SquiggleNet. Using the full length of the input sequence increased the accuracy of Guppy+Minimap2 to 91%, but the processing time increased dramatically. With UNCALLED, 32 threads were used to process 3000 signals, 6000 signals, and full-length reads respectively. It took at least 1277.28 seconds (21 min 17s) to finish the 3000 signals, but the accuracy was below 50%. With longer input length, the accuracy increased to 60% (82% for full-length), but the full length accuracy was still lower than SquiggleNet with 3000 signals. Meanwhile, the processing time grows drastically as well. SquiggleNet is, therefore, faster and more accurate than either Guppy+Minimap2 or UNCALLED.

**Figure 6:**
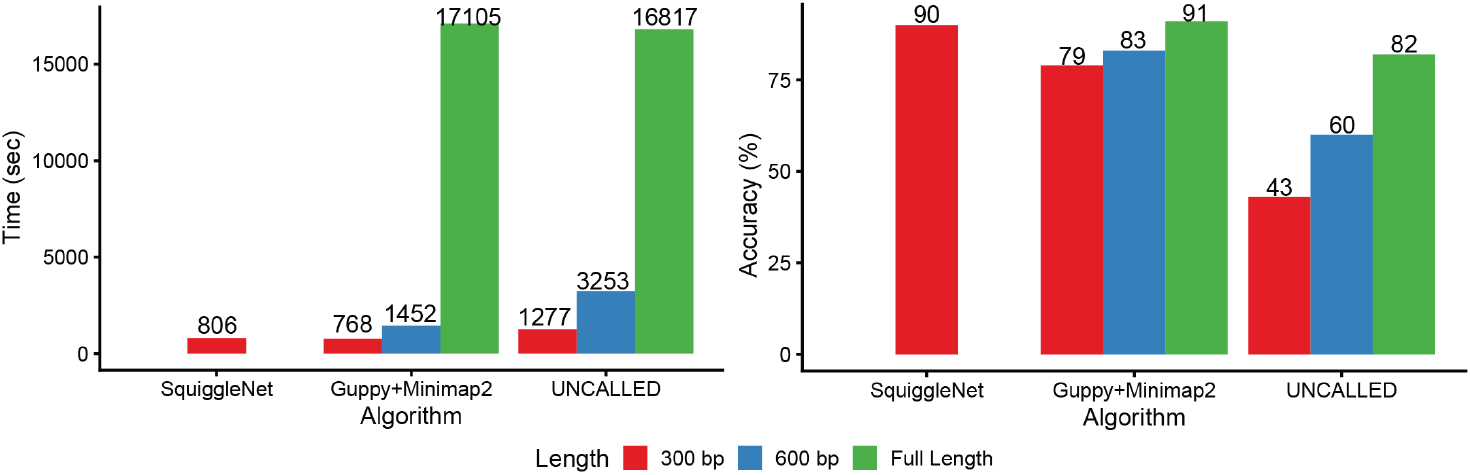
Processing Time and Accuracy Comparison: The processing time of SquiggleNet with 300 bp of input is among the lowest, and yet the accuracy is the highest among the three methods. For the other two alignment-based methods, with longer input length, the processing time grows drastically, whereas the accuracy gain is limited.

Note that the accuracy values that we report take into account all reads, including those that Minimap2 could not align. In contrast, the UNCALLED paper reported that the method was able to recover 94% of the alignments identified by Minimap2, but this number does not take into account the reads that Minimap2 failed to align. Crucially, our datasets have molecular barcodes, allowing us to determine the true species even for reads that Minimap2 failed to align. Furthermore, the 94% accuracy reported by the UNCALLED paper is based on using the entire sequence, whereas we only used significantly less information (3000 signals) to classify the reads.

We also observed that over 90% of the SquiggleNet processing time on GPU and over 40% of the processing time on CPU is spent on loading data from the disk. The actual classification time for a batch of 500 reads is about 0.06 seconds on GPU and about 0.8 seconds on CPU (one thread). When streaming directly from the nanopore sequencer in real time, this loading time can be significantly reduced.

SquiggleNet also offers significant advantages in terms of space requirements, requiring only 304KB to store the model parameters. The run-time space usage is dominated by the storage required for each mini-batch of sequences, rather than the model parameters. Guppy, however, is a much larger deep-learning model, and the smallest pre-trained option available through Oxford Nanopore Community [20] is 5.5MB. On top of that, however, the Guppy+Minimap2 method also requires a customized database for Minimap2 reference. In this experiment, the human and Zymo reference database takes 3.2GB. UNCALLED is currently operational only on CPU. Similarly, it also takes a reference database to build a Burrows-Wheeler index, which is an extra 3.2GB in this experiment. Therefore, SquiggleNet requires much less space than the other two methods.

### SquiggleNet Identifies Reads Containing Human Long Interspersed Repeat Elements

Classifying species is useful for many applications, but distinguishing among different loci from the same species would significantly expand the settings in which SquiggleNet can be applied. To investigate whether SquiggleNet can be used to enrich loci of interest within a single species, we analyzed data from a recently published protocol [21] that enriches specific families of interspersed repeat elements using Cas9-directed adapter ligation (Fig. 7a). This experimental enrichment is effective, but still fewer than 50% of reads contain the repeats of interest [21]. As previously described [22], we used BLASTn [23] on the base-called sequence of each read to label the reads as target (repeat-containing) or non-target (no repeat). We focused specifically on a single class of interspersed repeats, human-specific long interspersed elements (L1Hs), which was the family most effectively enriched by the Cas9-directed ligation protocol. Using these labels, we trained SquiggleNet on a balanced dataset containing approximately 170,000 reads from each class. Note that this is about tenfold less training data than we used in the human vs. bacterial classification experiments above. As with the species classification experiments, we discarded the first 1500 signals from each read, then used the next 3000 for training or testing. Despite the smaller training dataset, we found that SquiggleNet was extremely effective at identifying reads containing L1Hs elements (Fig. 7b-c). Our model achieved more than 92% accuracy with a true positive rate above 93%. These results indicate that using SquiggleNet in a read-until setting to enrich long interspersed repeats would provide a significant benefit compared to Cas9 enrichment only.

**Figure 7:**
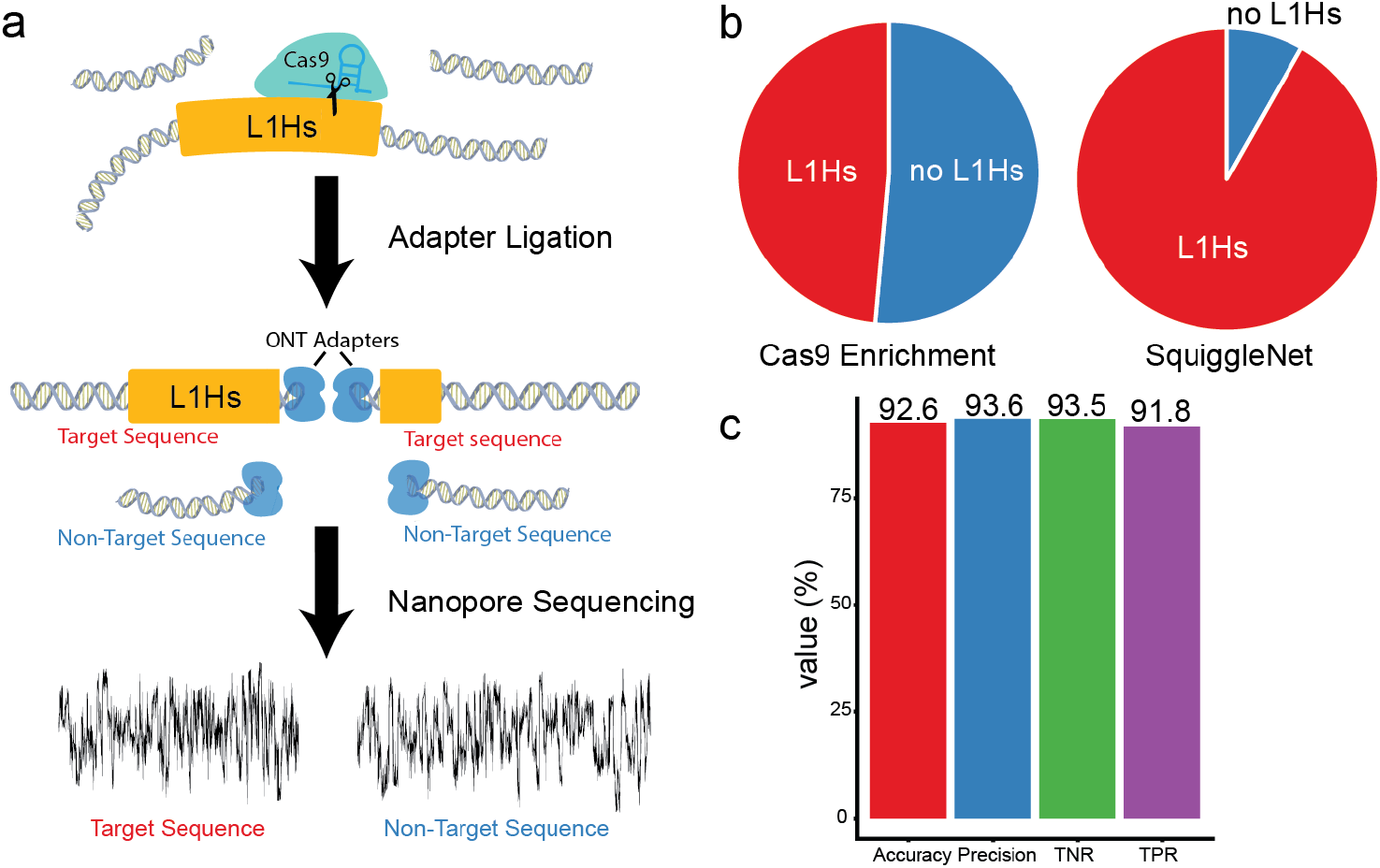
Identifying Reads Containing Human Long Interspersed Repeat Elements: (a) Diagram of experimental strategy for enriching human mobile elements, including interspersed repeats. A guide RNA specific to each repeat class directs Cas9 to cut the DNA and ligate a sequencing adapter. However, adapters are also ligated to some sequences without repeat elements. Subsequent nanopore sequencing produces both target and non-target reads. (b) Pie charts of the proportion of L1Hs repeat elements from Cas9 enrichment only vs. SquiggleNet classification. (c) Classification metrics demonstrating SquiggleNet’s ability to distinguish reads with or without L1Hs repeats.

### SquiggleNet Improves Throughput by Enabling Computationally Targeted Sequencing

To assess the potential improvement in sequencing throughput that SquiggleNet could provide, we developed a mathematical model to compare the total number of base pairs and total sequencing time needed to obtain a certain number of targeted sequences with and without SquiggleNet.

The detailed derivation of the model can be found in the Supplement. The most influential hyper-parameters include average target sequence length 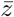, average non-target sequence length 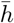, and target sequence concentration *c*. Several other tunable hyper-parameters, including the waiting time to eject one molecule and begin sequencing another; the total number of active pores in a flow cell; the sequencing speed; and the total number of targeted sequences, did not significantly influence the predicted increase in throughput (see Supplement). We chose the values for these less influential parameters based on the empirical time requirements and accuracy of SquiggleNet and the sample means from the real sequencing data.

In Figure 8, we picked the average non-target/target sequence length ratio as one axis, and the target sequence concentration as the other axis to demonstrate the total number of base pairs (left) and total sequencing time (right) that a regular nanopore sequencing pipeline would require, compared to those of a pipeline with SquiggleNet, in order to obtain a fixed number of targeted reads. Based on the properties of our sequencing datasets, we set average sequencing speed to be 450 base pairs per second and the total number of active pores in a flow cell to be 500. We also used the following parameters based on SquiggleNet’s empirical performance: TPR = 0.9, TNR = 0.9, sequencing time = 1s, and classification decision time = 0.8s (time required for SquiggleNet on CPU).

**Figure 8:**
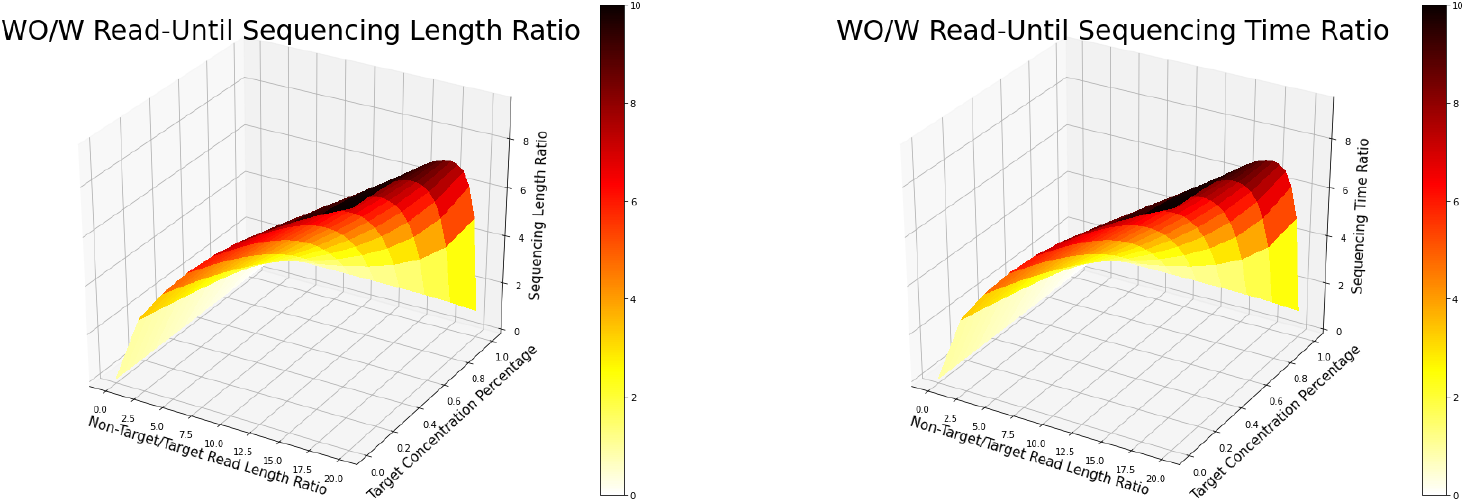
Throughput and Sequencing Time Comparison Without/With Read-Until: When the average non-target read length is about 20 times longer than the target read length, and sample contains over 90% non-target reads, a normal sequencing pipeline would have to sequence ∼10 times more base pairs (left) than Read-Until pipeline with SquiggleNet to achieve a fixed number of targeted reads. The ratio is about 10 for the required sequencing time as well (right).

We show the predicted gains in throughput and sequencing time for a range of the most important hyperparameters (Figure 8). When the average non-target read length is about 20 times longer than the target read length, and the sample contains over 90% non-target sequences, it would take a nanopore sequencing pipeline ∼10 times longer than Read-Until pipeline with SquiggleNet to achieve a fixed number of targeted reads. The regular nanopore sequencing pipeline would also have to sequence ∼10 times more base pairs than the Read-Until pipeline. Even if we set these parameters much more pessimistically, the model still predicts about a 5-fold gain in throughput and time. These numbers are also in the same ballpark as the 4.5× enrichment reported in the UNCALLED paper [10], supporting the plausibility of our mathematical model. We therefore conclude that Read-Until with SquiggleNet holds great promise to improve target read throughput, saving sequencing time and resources.

## Discussion

The success of our approach suggests that the raw sequencing signals generated by nanopore sequencing contain rich information for identifying target sequences from background sequences. Such features could include different DNA modification patterns, codon frequencies, GC content, or even DNA shape or RNA secondary structure. Furthermore, because these features are primarily local in nature, only a small amount of sequencing signal is required. In contrast, approaches that rely on sequence information alone require much more signal (more sequencing time), are susceptible to base-calling errors, and do not leverage non-sequence information. We also note that different reads go through the pores at different speeds. Future work could also include an event detector and a scaler into the classifier, which may further improve performance. Additionally, the squiggles from different MinION flow cells show systematic run-to-run differences. Thus, the data preprocessing and normalization procedures that we employed are crucial for generalizing across datasets.

We tested the capability of our model to enrich bacterial DNA in the presence of more abundant human DNA. Human and bacterial DNA are significantly different, which makes this classification task feasible. We also demonstrated that SquiggleNet can identify target sequences in other contexts, such as interspersed repeat elements within the human genome. This suggests that SquiggleNet could be used for targeted enrichment of other genomic loci, such as commonly mutated cancer genes or regions that are highly polymorphic across the human population.We also anticipate that SquiggleNet will be useful for distinguishing viral DNA or RNA sequences from host molecules within infected cells. Rapidly identifying targeted sequences could be helpful in numerous clinical settings, including cancer diagnosis, respiratory pathogen identification, and Coronavirus testing.

## Methods

### Data Collection

We generated five datasets using a MinION sequencer (Table 2) for this paper. The first four datasets used the standard Rapid Sequencing Kit (SQK-RAD004) protocol on a FLO-106D MinION Flow Cell. The HeLa&Zymo dataset used the ZymoBIOMICS Microbial Community DNA Standard, and Datasets 2−4 used the ZymoBIOMICS HMW DNA Standard with different barcodes specified in Table 2. Details about the dataset composition can be found in the Supplement. The Respiratory Metagenome data collection method can be found in [17].

**Table 1:** Method Requirement Comparison

|  | SquiggleNet | SquiggleNet | Guppy+Minimap2 | UNCALLED |
| --- | --- | --- | --- | --- |
| Equipment($t$=thread) | GPU | CPU($t$=1) | GPU+CPU($t$=32) | CPU($t$=32) |
| Space Requirement↓ | ≤ GPU Max | ≤ Mem Max | ≤ GPU Max + 3.2GB | ≤ Mem Max + 3.2GB |
| Model Size↓ | 304 KB | 304 KB | 5.5/40/116 MB |  |

**Table 2:** Datasets Description

| Name | Ratio | Train | Validation | Test | Note |
| --- | --- | --- | --- | --- | --- |
| HeLa&Zymo | 1:1 | 2.4M | 40k | 54k |  |
| Human&Zymo.b12 | 1:1 | 1.72M | 40k | 44k | Barcode 1 & 2 |
| Human&Zymo.b34 | 1:4 |  |  | 224k | Barcode 3 & 4 |
| Human&Zymo.b56 | 99:1 |  |  | 100k | Barcode 5 & 6 |
| Respiratory Metagenome | 951:1 |  |  | 324867 | Real Patient Samples |

Because base calling and sequence alignment of noisy nanopore reads can result in systematic errors and is not a completely reliable source of ground truth, we used barcodes to label the sequences in the three Human&Zymo datasets as either bacteria or human before mixing them together. The labels obtained in this way thus represent reliable ground truth.

Each extracted signal read was normalized with fast5 scaling and offset. All reads were also normalized using Z-scored median absolute deviation. The extreme signal values with a modified z-score larger than 3.5 were replaced by the average of closest neighbors.

### Model Architecture

SquiggleNet is a 1D-ResNet-based binary classifier (Figure 1). The first layer of 1D-CNN is comparable to the first layer of Guppy[20], but with significantly fewer channels (20 instead of 512). After that, there are four layers of 1D-ResNet, and each layer includes two BottleNeck blocks. The number of channels for each layer increases by a factor of 1.5, and each BottleNeck block decreases the string size with a stride of 2. We perform average pooling after the final convolutional layer, followed by a fully connected layer. We also experimented with other architectures (see Supplement).

### Training and Evaluation

Our best-performing model was trained on the HeLa&Zymo dataset with binary cross-entropy loss. The dataset was split into training, validation, and testing sets. The Human&Zymo b12, Human&Zymo b34, Human&Zymo b56, and Respiratory Metagenome datasets were used to assemble testing sets for the best-performing model.

As a separate analysis, we also trained on the Human&Zymo datasets and tested on the HeLa&Zymo datasets. The performance of this model was nearly identical but slightly worse than the model trained on HeLa&Zymo (see Supplement for details).

The Adam optimizer was used for over 6 epochs on each dataset, with a learning rate of 1e-3 and batch size of 1000. The model was initialized using Kaiming initialization in fan-out mode. Batch normalization was conducted within each Bottleneck block.

For the human interspersed repeat analysis, we used a balanced training dataset with equal numbers of reads that contained and did not contain an L1Hs element (about 170,000 reads each). We identified L1Hs elements using BLAST on the base-called reads with an e-value cutoff of 1 × 10^−5^, as previously described [21]. We trained for 3 epochs using the Adam optimizer. Evaluation metrics include overall accuracy, true positive rate (TPR, Recall), true negative rate (TNR), Precision, and area under receiver operating characteristic curve (AUROC) when running the model on different test datasets. Speed and memory comparisons were performed on the same Intel(R) Xeon(R) CPU E5-2697 v3 @ 2.60GHz machine with a single TITAN Xp GPU.

## Supplement

### Data Collection

We generated five datasets using a MinION sequencer (Table 2) for this paper. The first four datasets used the standard Rapid Sequencing Kit (SQK-RAD004) protocol on FLO-106D MinION Flow Cell. The HeLa&Zymo dataset used the ZymoBIOMICS Microbial Community DNA Standard, and Datasets 2−4 used the ZymoBIOMICS HMW DNA Standard with different barcodes specified in Table 2. The Respiratory Metagenome data collection method is described in [17].

The theoretical composition of the ZymoBIOMICS Microbial Community DNA Standard [12] includes 8 types of bacteria with 12% each of: *Listeria monocytogenes* (Lis), *Pseudomonas aeruginosa* (Pse), *Bacillus subtilis* (Bac), *Escherichia coli* (Esc), *Salmonella enterica* (Sal), *Lactobacillus fermentum* (Lac), *Enterococcus faecalis* (Ent), *Staphylococcus aureus* (Sta); and 2 types of fungi (2% each of *Saccharomyces cerevisiae* and *Cryptococcus neoformans*). The theoretical composition of the ZymoBIOMICS HMW DNA Standard [14] includes 7 types of bacteria with 14% each of *Listeria monocytogenes, Pseudomonas aeruginosa, Bacillus subtilis, Escherichia coli, Salmonella enterica, Enterococcus faecalis, Staphylococcus aureus* ; and 1 type of fungus (*Saccharomyces cerevisiae*, 2%).

The HeLa&Zymo dataset contains 200ng of HeLa DNA and 200ng of Zymo-BIOMICS Microbial Community DNA Standard computed by volume and listed concentrations. The sequenced samples were basecalled using Guppy v3.6.1 and aligned with reference downloaded from NCBI using Minimap2. We assigned the species of these sequences using the Minimap2 alignment because no species barcodes were used in this dataset. Due to the limited fungus sequences, only the bacterial sequences were kept for training, validation, and testing. The first 1500 signals for each read (about 150bp of sequence on average) were removed to avoid adapter sequences and signal instability.

The assembly of the Human&Zymo datasets each started with 400ng of Human GM12878 DNA and 400ng of ZymoBIOMICS HMW DNA Standard. They were barcoded using the SQK-RBK004 barcodes 1 and 2, 3 and 4, 5 and 6 respectively as indicated in the table. Then each dataset was generated using 200 ng of barcoded GM12878 (by volume) and 200ng of HMW Zymo (by volume) with a total of 400ng pooled in 10ul. For the Human&Zymo datasets, the extracted samples were basecalled using Guppy v3.6.1 with “–barcode kits SQK-RBK004” specification to separate the reads and identify the barcodes. The barcodes were then used to identify the species of each read. The first 2000 (about 200bp worth of sequences on average) signals were removed to avoid barcode overfitting and signal instability. Based on the adapter, barcode and barcode flanking sequence description from the Nanopore Community [24, 25], 2000 signals should be more than enough to remove all above.

Each extracted signal read was normalized with fast5 scaling and offset. All reads were also normalized using Z-scored median absolute deviation. The extreme signal values with a modified z-score larger than 3.5 were replaced by the average of closest neighbors.

### Model Architecture Experiments and Hyperparameter Tuning

In this section, we describe some of the additional models and hyperparameter settings that we explored while developing SquiggleNet. Table 3 summarizes these results. Most of the model architectures were trained on the HeLa&Zymo dataset using 1000 signals per sequence, after removing the first 1500 signals. We used this shorter input length and a subset of the training dataset to enable more rapid model testing. We then re-trained some of the more promising models with 3000 signals per read. Each model was trained for 2 to 5 epochs until the validation accuracy started to plateau or show signs of overfitting.

**Table 3:**
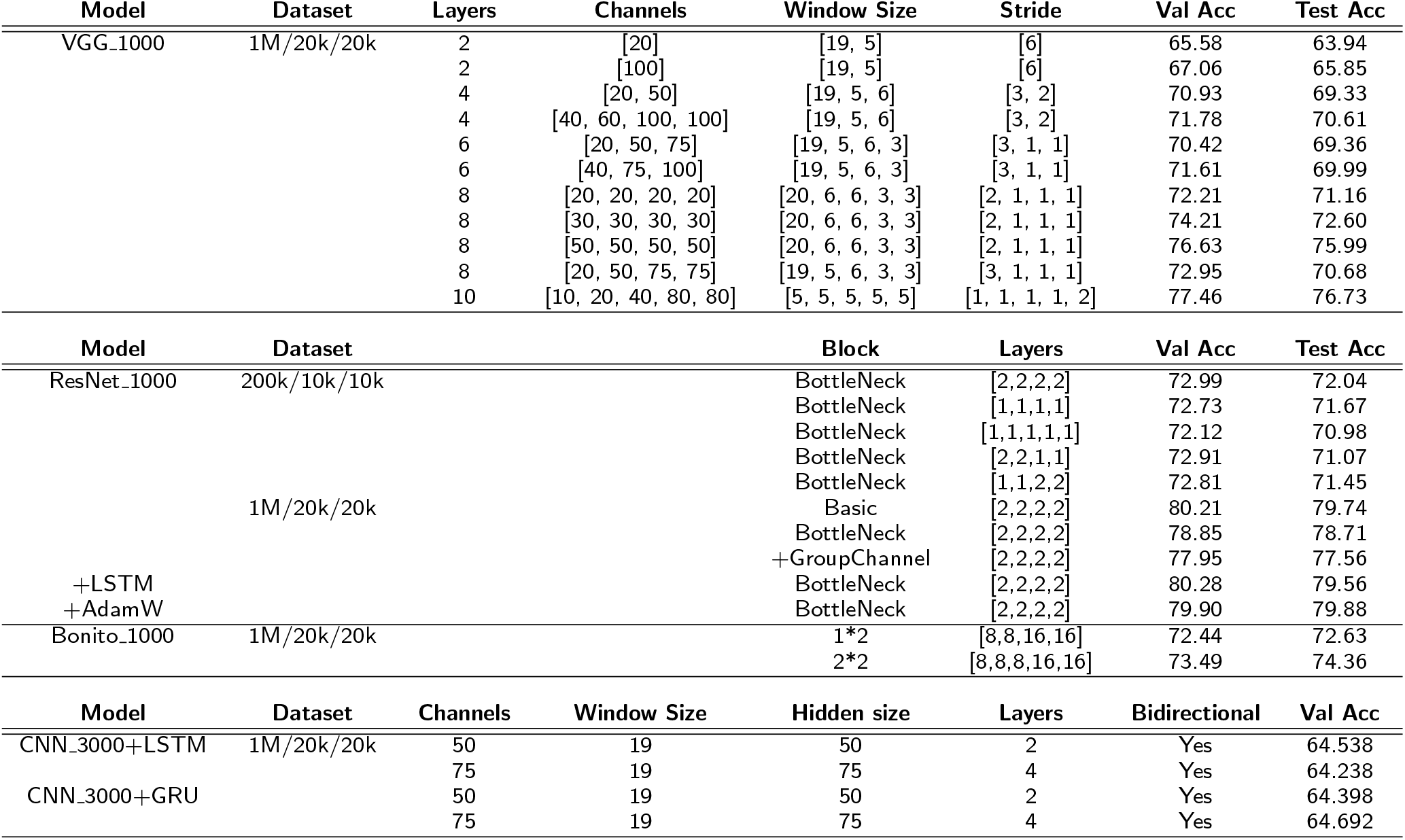
Selected Model Architecture Experiment Results and Hyperparameter Tuning Performance Report

The experiments based on the VGG 1000 architecture indicated that a model with more layers and more channels could generally offer better performance. However, the lack of shortcut skips in the architecture limited the possible depth of the VGG models.

We thus next experimented with a ResNet model with different numbers of layers, different numbers of blocks in each layer, different block types, and several other modifications. We found that increasing the number of blocks in each layer boosted performance somewhat, but increasing the number of layers did not necessarily improve the performance. Increasing the size of the training dataset boosted the model performance by ∼8% according to Table 3. We introduced a layer of LSTM in the middle of our ResNet model, and this increased performance by a small margin, but took much longer to train.

Recurrent Neural Network (RNN) models such as LSTM and GRU gave little advantage over CNN based models and also required significantly more resources to train. For the last set of experiments presented in Table 3, we used 3000 signals per read, introduced one layer of CNN with window size 19 to reduce the input size, and followed with various RNN architectures. The performance slowly increased from random guessing (50%), but plateaued around 65%. This could indicate that local features extracted by convolution provide sufficient information for classification, and long-range dependencies extracted by the recurrent network only help by a small amount.

Other models we tried include: a down-sized Bonito [26, 27], stacks of LSTM layers with variational window sizes, different hyperparameter settings, and different training datasets. After full consideration of model size, speed, performance, and training time, we settled on the best performing model architecture to perform the main experiments in the paper.

### Comparison of Models Trained on HeLa&Zymo and Human&Zymo

In addition to the best-performing model trained on the HeLa&Zymo dataset, we conducted the same set of test experiments on a model trained on the Human&Zymo b12 dataset. This dataset consists of a 1:1 mix of human DNA and Zymo HMW DNA, as described in Supplement Section Data Collection. We tested on all five datasets, which include different sample preparations, flow cells, sample components, and human:bacteria ratios. The performance can be found in Figure 9. The overall performance is almost identical with Figure 2, but slightly worse. This may be because the Hyman&Zymo b12 dataset contains less training data than the HeLa&Zymo dataset. Overall, this analysis indicates that the model achieves high accuracy whether trained on HeLa DNA (which is a highly mutated cancer cell line) or GM12878 DNA (which should look more like healthy human DNA), when tested on the other type of data.

**Figure 9:**
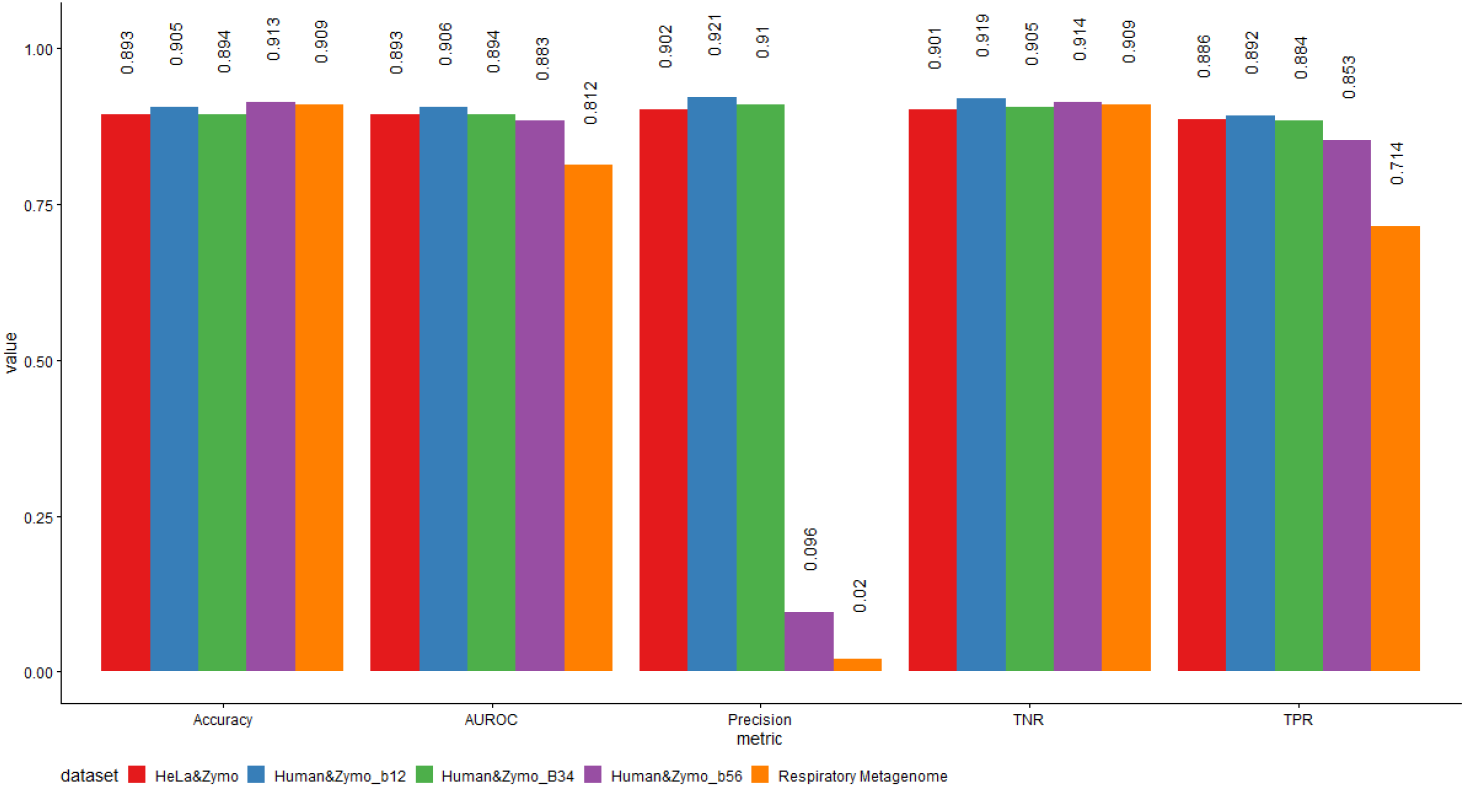
Overall Performance across five test Datasets with Model trained on Hyman&Zymo b12: Accuracy, True Positive Rate (TPR, RECALL), True Negative Rate (TNR), Precision, and the AUROC score of the model trained on the HeLa&Zymo training set, and tested on five test sets with bacterial sequences as the target.

### Comparison Experiment Method Details

The performance and efficiency comparison experiment was conducted on dataset Human&Zymo b34 with 1:4 Human and Zymo mix. For a fair read-in and write-out time comparison across the platforms, all the reads were pre-truncated into 3000 signals, 6000 signals, and full length, after removing the first 2000 signals for adapters and barcode. Any reads shorter than the minimum input requirements were discarded. The ground truth labels for each read were obtained by the SQK-RBK004 barcode 3 and 4, basecalled by Guppy v3.6.1. Any reads that were labeled as other barcodes or not labeled were discarded.

SquiggleNet was tested on 1) a single-usage Intel(R) Xeon(R) CPU E5-2697v3 @ 2.60GHz machine with a single TITAN Xp GPU with batch size one (fast5 file, about 4000 sequences each), and 2) a single-usage 3.5 GHz Dual-Core Intel Core i7 Macbook Pro with a single thread. Over 90% of the compute time on GPU was spent on file read-in and write-out (over 40% of the time on CPU).

When testing the Guppy+Minimap2 method, we used 4 callers and 4 runners per GPU for Guppy, and 32 threads for Minimap2 sequence alignment. Indexes for both human and Zymo community reference genome were pre-generated. The test datasets with 3000 signals per read, 6000 signals per read, and full length reads were basecalled by Guppy and then aligned using Minimap2 with the pre-generated index. The resulting read IDs were cross-referenced with the ground truth labels for barcode 3 and 4 to calculate the overall accuracy.

When testing the UNCALLED method, we used 32 threads to process 3000 signals, 6000 signals, and full-length reads. BWA index was pre-generated for the Zymo community reference genome. Unfortunately, UNCALLED struggles to map repetitive references longer than 100Mbp, such as the human genome. Therefore, we had to treat all the reads that weren’t classified as Zymo to be human reads as an approximation for the performance computation. The resulting read IDs were cross referenced with the ground truth labels for barcode 3 and 4, to calculate the overall accuracy.

### DNA Methylation Differences Between Human and Bacterial DNA

Since SquiggleNet achieves higher classification accuracy than base calling followed by sequence alignment (given 3000 signals), we hypothesized that the neural network may be using other information besides just nucleotide sequence. One possible such source of information is methylation. We therefore investigated whether there are systematic methylation differences between the human and bacterial DNA sequences.

To do this, we used Guppy v3.6.1 to output the methylation probability for each sequence in Dataset Hyman&Zymo b34. For each base pair with greater than zero percent chance to be an A or C (the only nucleotides for which Guppy will predict methylation), we computed the most likely base call: A, methylated A, C, or methylated C. We then calculated the total and average numbers of methylated As and methylated Cs in the first 1000 bases of both human and bacterial sequences.

As shown in Table 4, bacterial sequences contain on average about 2 methylated A nucleotides and 4 methylated C nucleotides per kilobase, whereas the human sequences contain on average 0 methylated As and 16 methylated Cs per kilobase. This suggests that methylation is indeed a possible feature that could help to classify sequences as bacterial or human.

**Table 4:** Average number of methylated A (mA) and methylated C (mC) nucleotides in the first 1000 nucleotides of human and bacterial DNA sequences

|  | Avg mAs/KB | Avg mCs/KB |
| --- | --- | --- |
| Bacterial | 1.72 | 3.53 |
| Human | 0.01 | 16.27 |

### Model Interpretation Using Integrated Gradients

We used the method of integrated gradients (IG) [15] to investigate the relationship between convolutional filters, sequencing signal dynamics, and classification results from SquiggleNet. Integrated gradients computes the amount of gradient change for each corresponding input, and by doing so, offers interpretation on which part of the input contributes the most to the model’s decision. The IG approach gives a way of attributing model outputs to specific input features. In our case, IG can tell us which electrical signal values over the course of the sequencing time most strongly influence SquiggleNet’s decision to classify the sequence as positive or negative.

To investigate this, we adapted an existing PyTorch implementation [28] of IG so that it works for 1-D squiggles, rather than 2-D images. We used a squiggle whose value is identically the mean pore conductance (which is identically 0 after normalization) as the background signal required by IG. This is analogous to using an all-black image (the standard background signal for performing IG on image data). Then we simultaneously plotted the raw squiggle, base calls, and integrated gradients. Figure 10 shows the results for two positive and two negative sequences, with one positive and one negative containing a methylated nucleotide. The attribution is clearly strongest at positions where the signal changes direction and/or changes by a large amount. Attribution also tends to be high at the nucleotide boundaries predicted by the base caller. This suggests that SquiggleNet has learned filters related to the nucleotide composition of the signal and uses the results to make classification decisions.

**Figure 10:**
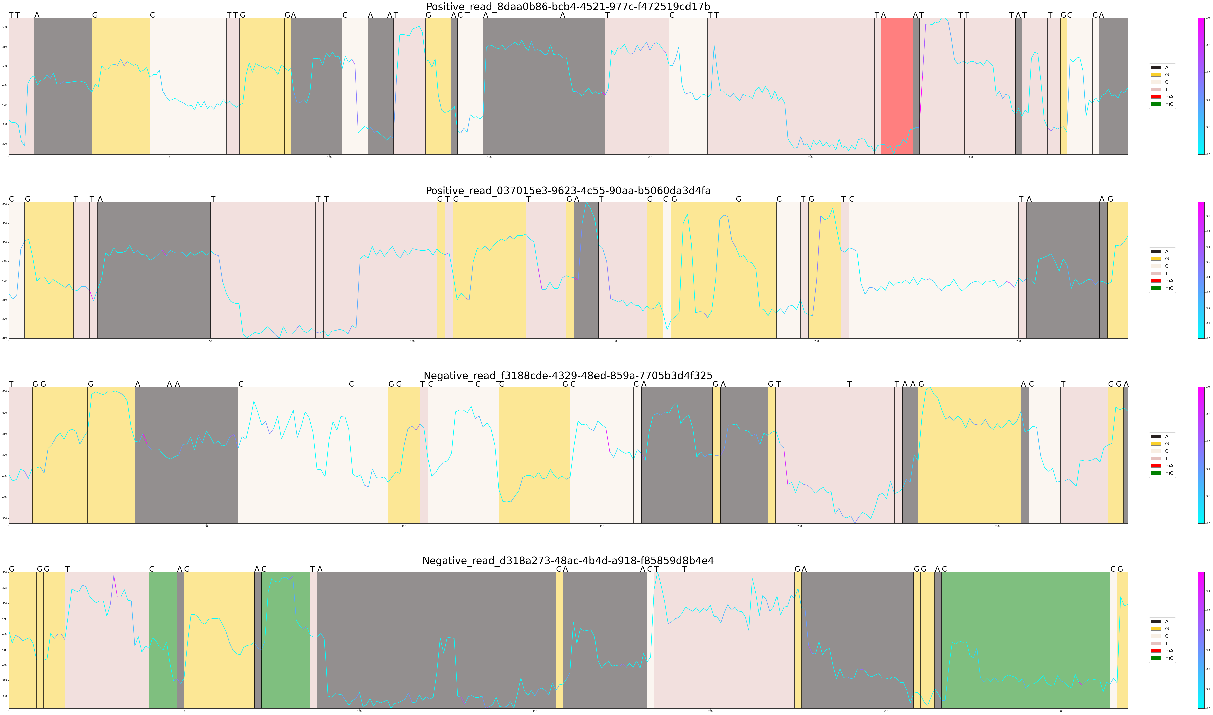
SquiggleNet Integrated Gradients and Signal Dynamics. Four sample signal chunks, 2 positive signals (top), and 2 negative signals (bottom). Each signal chunk was colored with corresponding predicted nucleotide (from Guppy basecalling results). Methylated As were colored in red, and methylated Cs were colored in green. Each squiggle was also colored by the value of the integrated gradient. A higher integrated gradient value (purple) indicates that the model is more excited by that part of the signal, and thus that it contributes more to the final classification decision.

### Throughput Estimation Model

The throughput estimation model computes the total amount of time *t* and the number of base pairs *l* needed to achieve *x* targeted reads, and compares these values with and without read-until.

The estimation model assumes the average targeted read length to be 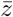, and the average non-targeted read length to be 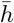. It assumes the total number of functioning pores *k* is constant throughout the sequencing run. It also assumes all the functioning pores are actively sequencing or in the process of getting a new read. All the other parameter values can be found in table 5.

**Table 5:** Throughput Estimation Model Parameters

| Description | Symbol | Value |
| --- | --- | --- |
| Total sequencing time | $t$ | |
| Total base pair sequenced | $l$ | |
| Number of targeted reads | $x$ | 1000 |
| Number of total reads | $n = x/a$ | |
| Number of pores alive | $k$ | 400 |
| Average sequencing speed | $v$ | 450 bp/s |
| Average read length: target | $\bar{z}$ | 40000 bp |
| Average read length: non-target | $\bar{h}$ | 40000 bp |
| Concentration: target | $a$ | 0.1 |
| Concentration: non-target | $b = 1 - a$ | 0.9 |
| Classifier TPR | $p_1$ | 90% |
| Classifier TNR | $p_2$ | 90% |
| Time: attract a new read | $t_0$ | 0.01s |
| Time: initial sequencing | $t_1$ | 1s |
| Time: decision time | $t_2$ | 0.8s |
| Time: reject a read | $t_3$ | 0.01s |

Without Read-Until, the total sequencing time *t*_*wo*_ and the total number of base pairs *l*_*wo*_ needed for *x* targeted sequences are:

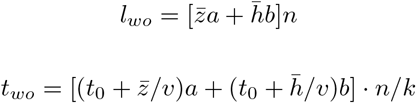

With Read-Until, the total sequencing time *t*_*ru*_ and the total number of base pairs *l*_*ru*_ needed for *x* targeted sequences are:

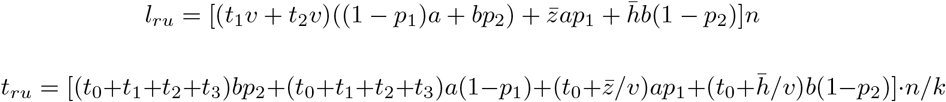

Several parameters cancel out and thus do not affect the sequencing length ratio *l*_*wo*_*/l*_*ru*_ or the sequencing time ratio *t*_*wo*_*/t*_*ru*_. These parameters include: the number of targeted reads *x*, the number of total reads *n*, and the number of active nanopores *k*. Several hyperparameters were set to default values based on SquiggleNet’s statistics and the sample means from multiple sequencing experiments. These include average sequencing speed (*v* = 450 bp/s), classification TPR (*p*_*1*_ = 90%), classification TNR (*p*_*2*_ = 90%), initial sequencing time (*t*_*1*_ = 1*s*), and decision time (*t*_*1*_ = 0.8). The average read length for target and non-target reads are both set to 40000 bp. We set the target concentration to 10% and non-target read concentration to 90%.

We also investigated how *t*_*0*_, the time to attract a new read, or *t*_*3*_, affect the throughput ratio. We varied these two parameters from 0 to 1 second and plotted the resulting throughput ratio for both sequencing length and sequencing time (Figure 11). Neither parameter significantly affected the throughput ratio.

**Figure 11:**
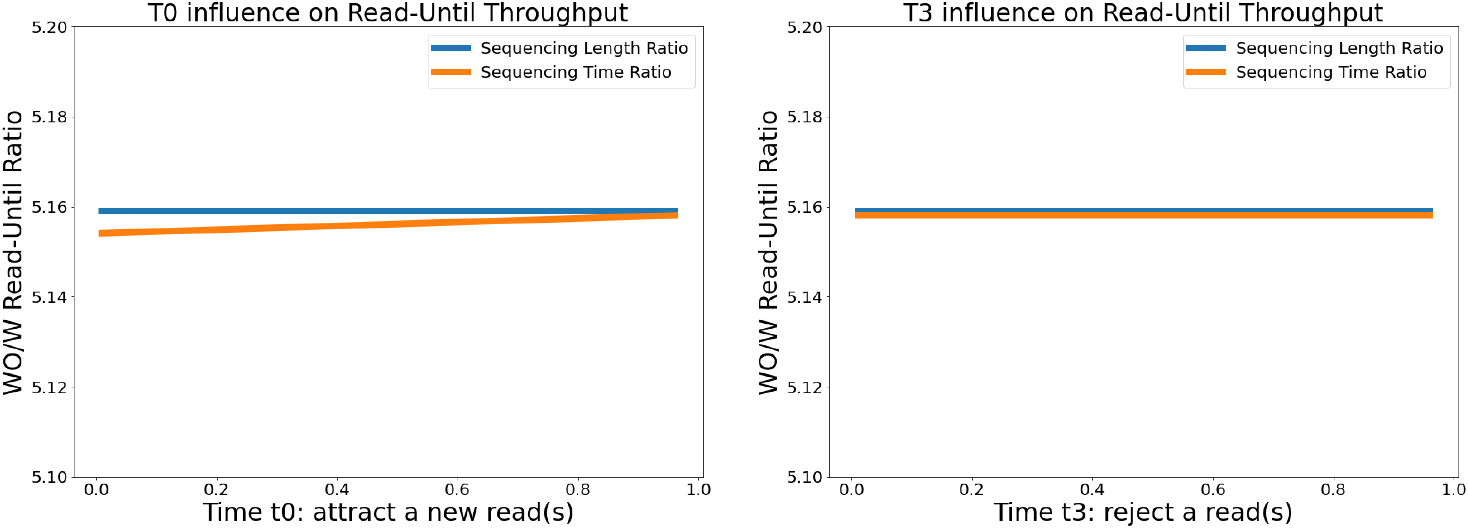
The influence of t0 and t3 on Read-Until Throughput: As t0 and t3 change from 0 to 1 second, the sequencing time and length ratio (without/with read-until) are not effected significantly.

### Model Performance on Different Test Dataset Composition Ratio

Because the trained model makes predictions independently for each individual sequence, the true positive, false positive, false negative, and true negative rates are independent of the distribution of the test set. In contrast, the ratio of sequences in the training dataset is important, which is why we conducted training with equal numbers of positive and negative sequences. To confirm this, we used the model pretrained on the HeLa&Zymo dataset (same as reported in Figure 2), and tested it on datasets with HeLa&Zymo composition ratios ranging from 100:0, 99:1, 19:1, 3:1, 1:1, to 1:3, 1:19, 1:99 to 0:100. As Figure 12 shows, the TPR/TNR/FNR/FPR values are not significantly affected by the test composition ratio and remain around 90%. The absolute number of positive and negative predictions will of course vary with the composition of the test dataset, but this behavior is desirable.

**Figure 12:**
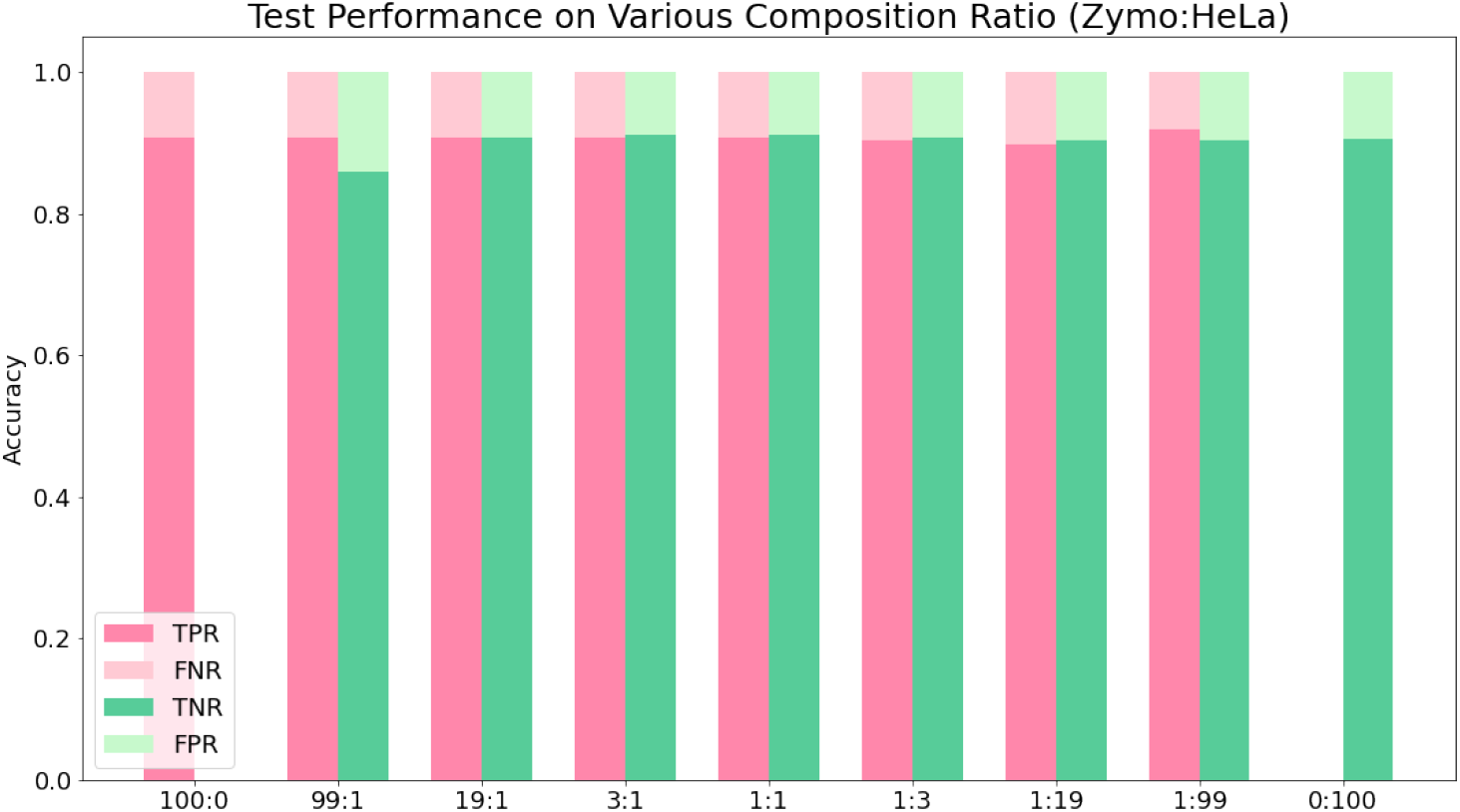
Robustness of SquiggleNet Performance Across Varying Test Set Composition: The TPR, FNR, TNR, and FPR are independent of the test dataset Zymo:HeLa ratio and remain around 90%.

### Model Performance on Odd and Even HeLa Chromosomes

We trained a new model on the HeLa & Zymo dataset using even chromosomes for training, and odd for testing, each with a 1:1 ratio of HeLa and Zymo sequences. We omitted all chromosomes with non-numeric names–including the mitochondrial chromosome, sex chromosomes, and other contigs–rather than arbitrarily designating them as odd or even. Otherwise we followed the training procedure exactly as described in the Methods. The analysis indicates that SquiggleNet can correctly classify reads from missing chromosomes (Table 6). The performance is about 6% higher than what we obtained using all chromosomes. This is possibly due to the chromosomes we excluded, which may be enriched for low-quality sequences. In addition, the human mitochondrial chromosome is somewhat phylogenetically similar to prokaryotic DNA, so excluding it may have also boosted the performance.

**Table 6:** Model Performance on Odd and Even HeLa Chromosomes

| Accuracy | Precision | TPR (Recall) | TNR |
| --- | --- | --- | --- |
| 96.84% | 96.97% | 96.65% | 97.02% |

## Availability of data and materials

We have uploaded our data to SRA (SRP296988). The Cas9 enrichment dataset is previously published and available on the SRA (BioProject accession PRJNA699027). The code can be found in our Github repository: https://github.com/welch-lab/SquiggleNet.

## Author’s contributions

YB and JDW conceived the idea of SquiggleNet. YB designed and implemented the model and analyzed the data. JW generated datasets 1 to 4. JRE, PR and RPD provided the human-pathogen dataset and generated Figure 5. YB and JDW wrote the paper. JRE and DB offered suggestions and advice. TM, WZ, RM, and AB generated and annotated the Cas9 enrichment dataset. All authors read and approved the final manuscript.

## Acknowledgements

We thank members of the CELab and Prof. Jenna Wiens for their helpful suggestions and advice during this project.

## References

1. Oxford Nanopore: Minion. https://nanoporetech.com/products/minion

2. Gnirke, A., Melnikov, A., Maguire, J., Rogov, P., LeProust, E.M., Brockman, W., Fennell, T., Giannoukos, G., Fisher, S., Russ, C., Gabriel, S., Jaffe, D.B., Lander, E.S., Nusbaum, C.: Solution hybrid selection with ultra-long oligonucleotides for massively parallel targeted sequencing. Nature Biotechnology 27(2), 182–189 (2009). doi:10.1038/nbt.1523

3. Kozarewa, I., Armisen, J., Gardner, A.F., Slatko, B.E., Hendrickson, C.L.: Overview of target enrichment strategies (1934-3647 (Electronic))

4. Rand, A.C., Jain, M., Eizenga, J.M., Musselman-Brown, A., Olsen, H.E., Akeson, M., Paten, B.: Mapping dna methylation with high-throughput nanopore sequencing. Nature Methods 14(4), 411–413 (2017). doi:10.1038/nmeth.4189

5. Simpson, J.T., Workman, R.E., Zuzarte, P.C., David, M., Dursi, L.J., Timp, W.: Detecting dna cytosine methylation using nanopore sequencing. Nature Methods 14(4), 407–410 (2017). doi:10.1038/nmeth.4184

6. Charalampous, T., Kay, G.L., Richardson, H., Aydin, A., Baldan, R., Jeanes, C., Rae, D., Grundy, S., Turner, D.J., Wain, J., Leggett, R.M., Livermore, D.M., O’Grady, J.: Nanopore metagenomics enables rapid clinical diagnosis of bacterial lower respiratory infection. Nature Biotechnology 37(7), 783–792 (2019). doi:10.1038/s41587-019-0156-5

7. Gilpatrick, T., Lee, I., Graham, J.E., Raimondeau, E., Bowen, R., Heron, A., Sedlazeck, F.J., Timp, W.: Targeted nanopore sequencing with cas9 for studies of methylation, structural variants, and mutations. bioRxiv, 604173 (2019). doi:10.1101/604173

8. Gu, W., Crawford, E.D., O’Donovan, B.D., Wilson, M.R., Chow, E.D., Retallack, H., DeRisi, J.L.: Depletion of abundant sequences by hybridization (dash): using cas9 to remove unwanted high-abundance species in sequencing libraries and molecular counting applications. Genome Biology 17(1), 41 (2016). doi:10.1186/s13059-016-0904-5

9. Payne, A., Holmes, N., Clarke, T., Munro, R., Debebe, B.J., Loose, M.: Readfish Enables Targeted Nanopore Sequencing of Gigabase-sized Genomes. doi:10.1038/s41587-020-00746-x

10. Kovaka, S., Fan, Y., Ni, B., Timp, W., Schatz, M.C.: Targeted Nanopore Sequencing by Real-time Mapping of Raw Electrical Signal with UNCALLED. doi:10.1038/s41587-020-0731-9

11. He, K., Zhang, X., Ren, S., Sun, J.: Deep Residual Learning for Image Recognition (2015). 1512.03385

12. ZymoBIOMICS Microbial Community DNA Standard. https://www.zymoresearch.com/collections/zymobiomics-microbial-community-standards/products/zymobiomics-microbial-community-dna-standard

13. Li, H.: Minimap2: pairwise alignment for nucleotide sequences. Bioinformatics 34(18), 3094–3100 (2018). doi:10.1093/bioinformatics/bty191. https://academic.oup.com/bioinformatics/article-pdf/34/18/3094/25731859/bty191.pdf

14. ZymoBIOMICS HMW DNA Standard. https://www.zymoresearch.com/collections/zymobiomics-microbial-community-standards/products/zymobiomics-hmw-dna-standard

15. Sundararajan, M., Taly, A., Yan, Q.: Axiomatic attribution for deep networks. In: Precup, D., Teh, Y.W. (eds.) Proceedings of the 34th International Conference on Machine Learning. Proceedings of Machine Learning Research, vol. 70, pp. 3319–3328. PMLR, International Convention Centre, Sydney, Australia (2017). http://proceedings.mlr.press/v70/sundararajan17a.html

16. Oxford Nanopore Technologies, M.L.: Real-Time Selective Sequencing on the MinION. Youtube. https://www.youtube.com/watch?v=34sWScdYyYQ&t=303s&ab_channel=OxfordNanoporeTechnologies

17. O’Dwyer, D.N., Ashley, S.L., Gurczynski, S.J., Xia, M., Wilke, C., Falkowski, N.R., Norman, K.C., Arnold, K.B., Huffnagle, G.B., Salisbury, M.L., Han, M.K., Flaherty, K.R., White, E.S., Martinez, F.J., Erb-Downward, J.R., Murray, S., Moore, B.B., Dickson, R.P.: Lung microbiota contribute to pulmonary inflammation and disease progression in pulmonary fibrosis. American journal of respiratory and critical care medicine 199(9), 1127–1138 (2019). doi:10.1164/rccm.201809-1650OC

18. Pendleton, K.M., Erb-Downward, J.R., Bao, Y., Branton, W.R., Falkowski, N.R., Newton, D.A.-O., Huffnagle, G.A.-O., Dickson, R.A.-O.: Rapid pathogen identification in bacterial pneumonia using real-time metagenomics (1535-4970 (Electronic))

19. Ondov, B.D., Bergman, N.H., Phillippy, A.M.: Interactive metagenomic visualization in a web browser. BMC Bioinformatics 12(385), 1471–2105 (2011)

20. Oxford Nanopore: Guppy. https://community.nanoporetech.com/protocols/Guppy-protocol/v/GPB_2003_v1_revT_14Dec2018

21. McDonald, T.L., Zhou, W., Castro, C.P., Mumm, C., Switzenberg, J.A., Mills, R.E., Boyle, A.P.: Cas9 targeted enrichment of mobile elements using nanopore sequencing. Nature Communications 12(1), 3586 (2021). doi:10.1038/s41467-021-23918-y

22. Zhou, W., Emery, S.B., Flasch, D.A., Wang, Y., Kwan, J.M.K.Y., Kidd Moran, J.V., Mills, R.E.: Identification and characterization of occult human-specific line-1 insertions using long-read sequencing technology. Nucleic Acids Res 48(3), 1146–1163 (2020). doi:10.1093/nar/gkz1173

23. Zhang, Z., Schwartz, S., Wagner, L., Miller, W.: A greedy algorithm for aligning dna sequences. J Comput Biol 7(1-2), 203–14 (2000). doi:10.1089/10665270050081478

24. Oxford Nanopore: Barcoding Kits. https://community.nanoporetech.com/technical_documents/chemistry-technical-document/v/chtd_500_v1_revw_07jul2016/barcoding-kits

25. Oxford Nanopore: Rapid Sequencing Kit Family. https://community.nanoporetech.com/technical_documents/chemistry-technical-document/v/chtd_500_v1_revw_07jul2016/rapid-sequencing-kit-family

26. Kriman, S., Beliaev, S., Ginsburg, B., Huang, J., Kuchaiev, O., Lavrukhin, V., Leary, R., Li, J., Zhang, Y.: QuartzNet: Deep Automatic Speech Recognition with 1D Time-Channel Separable Convolutions (2019). 1910.10261

27. Bonito. https://github.com/nanoporetech/bonito

28. Integrated Gradient. https://github.com/TianhongDai/integrated-gradient-pytorch

